# Assessing mechanisms driving phenol isopropylated phosphate (IPP)-induced larval photomotor response deficits in zebrafish

**DOI:** 10.1101/2024.06.20.599969

**Authors:** Sunil Sharma, Alfredo Rojas, Rosemaria Serradimigni, Connor Leong, Subham Dasgupta

## Abstract

Phenol isopropylated phosphates (IPP) are an additive organophosphate flame retardant (OPFR) which has been extensively used in furniture, electronics, automobiles, plastics, and children’s products to slow down the spread of fire. The processing and distribution of IPP-containing products have been prohibited but its continuous leaching from end use products has retained the concern of its toxicity. The present study was designed to evaluate IPP-induced developmental toxicity using zebrafish embryos. We first conducted range finding experiments with embryonic zebrafish exposures to 0-200 μM IPP from 6 to 120 h post fertilization and found significant morphological impacts like pericardial edema, yolk sac edema and spinal curvature at higher concentrations. For behavioral readouts, we performed larval photomotor response (LPR) assay at sublethal concentrations and observed hypoactive locomotory behavior in exposed larvae. Following this, relying on secondary analyses of our whole embryo mRNA-seq data, we conducted-1) retinoic acid receptor (RAR) signaling assay and 2) DNA methylation assays. *In vitro* assay for RA receptors indicate that IPP significantly inhibits RARα, but not RARβ and RARγ. Whole-mount immunohistochemistry for 5-methylcytosine and global DNA methylation assay showed significant IPP-induced hypermethylation *in situ*. We conducted IPP co-exposure studies with a methylome modifier 5-azacytidine (Aza-c a methylation inhibitor) or retinoic acid signaling activators to assess if LPR phenotypes were mitigated by co-exposures. Data showed that Aza-c co-exposures partially reversed IPP-induced LPR hypoactivity and DNA hypermethylation, co-exposure with retinoic acid as well as AM580 (an RARα activator) were not able to reverse IPP-induced hypoactivity. Finally, based on RNA-seq data, we hypothesized that IPP affects the development of brain and eyes. Firstly, we performed global DNA methylation in brain and eyes, but did not find any significant effects. Then, we conducted mRNA sequencing on dissected brains and eyes, and found 2 and 135 differentially expressed genes, respectively. Gene ontology revealed that IPP affect phototransduction, voltage gated ion channels, synaptic and neurotransmitter signaling. Collectively, our data shows that IPP induces morphological abnormalities and disrupts larval photo motor response, potentially through methylomic regulation. Finally, we observed that IPP affects gene expression within the developing eye, establishing synaptic transmission, vision and muscle contraction as a potential causative factor for LPR responses.

## 1. Introduction

Over the past 20 years, there has been an increase in the use of organophosphate compounds as plasticizers and flame retardants (FRs) in consumer goods and construction materials (Wei et al., 2015; Fu et al., 2022). Organophosphate FRs (OPFRs) have gained popularity as suitable substitutes for brominated FRs and polychlorinated biphenyls (PCBs) which have been found persistent in the environment and toxic to animals (Dasgupta et al., 2018; Yao et al., 2021). OPFRs have been extensively used in every day-use product including household, textile, electronic, automobile and chemicals. Many of these are additive type of FRs which makes them prone to leach into the environment via volatilization, dissolution and attrition (Ekpe et al., 2020). These compounds have been detected in water (Xing et al., 2020), dust (Harrad et al., 2016), air (Han et al., 2020), sediments (He et al., 2017) and human beings (Guo et al., 2016). Because of hand to mouth exposure, observed concentration of OPFRs were higher in children i.e., 0.36 – 798 ng/ml compared to adults i.e., 0.39 – 104 ng/ml (Butt et al., 2016; Hoffman et al., 2015). OPFRs have been shown to cause developmental toxicity through epigenetic modifications (Volz et al., 2016), impaired neurodevelopment (Zhang et al., 2023), oxidative stress and endocrine disruption (Chen et al., 2015). Specifically, Castorina et al. (2017) found a significant correlation between increased exposure to specific OPFRs during pregnancy and childhood with neurodevelopmental disorders.

Phenol isopropylated phosphate (IPP) are an organophosphate isomeric FR mixture, with 2015 annual production was approximately 6 million lbs. It has been extensively used in a variety of products, such as rubber, polyurethane foam, textiles, casting resins, thermoplastics, electrical equipment, epoxy and phenolic resins to slow down the spread of fire. Although the processing and distribution of IPP containing products have been prohibited in 2021, but its isomers were detected in breast milk, urine and blood of humans, up to 100 ng/mL (Butt et al., 2014; Hoffman et al., 2017; Araki et al., 2018; Ospina et al., 2018). Triphenyl phosphate (TPHP), Mono-IPP, Bis-IPP and Tris-IPP are the major isomers of IPP with percent composition-21.5, 36.9, 21.9 and 8.5 %, respectively (Witchey et al., 2023). IPP and TPHP are two nominated OPFRs by the US consumer and product safety commission (CPSC) due to concern regarding increased hand to mouth exposure in children (Butt et al., 2016; Hoffman et al., 2017). Based on environmental monitoring, the predicted environmental concentration of TPHP was 25 ng/L, 293 ng/L, 9.1 μg/kg and 11 μg/kg in surface water, sewage treatment plant, soil and sediment, respectively (Australian Government, Evaluation statement, 2023). In Australia, TPHP levels have been found to be 51μg/kg in various indoor environments (Banks et al., 2020; Huang et al.,2020). Hydrophobic nature, easy adsorption, and stability makes it toxic and environmentally persistent (Mihajlovic et al., 2011). It has been reported that TPHP caused oxidative stress, cytotoxicity, metabolic disorders, endocrine disruption, neurodevelopmental toxicity and increased risk of cancers (Xie et al., 2023; Tachachartvanich et al., 2024). Wade et al. (2019) observed that isopropylated triphenyl phosphate (IPTPP) exposure to adult wistar rat results in hypertrophy and accumulation of lipids in zona fasciculate cells of adrenal cortex. Recent study of Witchey et al. (2023) reported that IPP and TPHP (1000 – 30000 ppm) exposure caused reproductive (decreased number of pups and delayed puberty) and developmental toxicity (altered organ and body weight) in Sprague Dawley rats. TPHP exposure triggered abnormal behaviour, visual impairment, apoptosis, oxidative stress and neurodevelopmental toxicity in early life stages of zebrafish. (Zhang et al., 2023). Due to ubiquitous nature, toxicity evidence and limited information, a thorough assessment of chemico-biological interactions of IPP during development is required to reduce risk, develop preventative and therapeutic strategies.

Our study specifically IPP developmental toxicity using zebrafish embryos as a model. Early life stages of zebrafish are vulnerable to the effects of environmental chemicals because of undergoing critical periods of development (Daston et al., 2004). Developing brain is sensitive to many xenobiotics which can alter the structure and function, resulting to adverse effects on learning, behavior and overall health (Rice and Barone, 2000). We exposed embryos at blastulation stage of embryogenesis to higher concentrations (20 and 200 μM IPP) and observed phenotypic anomalies. Following this, we also observed concentration-dependent hypoactive behavior in larval photomotor response (LPR)- a behavioral readout-during late embryogenesis. Since previous work by Dasgupta et al. (2021) suggested that IPP potentially affects neuronal methylation and retinoic acid signaling, major goal of this research was to study the mechanism of IPP induced hypoactive behavior. The major hypothesis were - 1) IPP induced hypoactive behavior is caused by a combination of DNA methylation and retinoic acid signaling and 2) IPP specifically affects eye and brain development.

## 2. Methods

### 2.1. Zebrafish husbandry

Wild type adult zebrafish (5D) were generously donated by Dr. Robyn Tanguay at the Sinnhuber Aquatic Research Laboratory and maintained at 28 ± 2 ^0^C temperature and 14h/10h, light/dark photoperiod on a recirculating system. The breeding and exposure to early life stages were performed with approved protocols (AUP2022-0434 and AUP2023-0114) provided by Institutional Animal Care and Use Committee (IACUC), Clemson University, South Carolina, USA.

### 2.2. Chemicals

IPP (CAS no 68937-41-7, Technical grade) was purchased from Toronto Research Chemicals; 5-azacytidine (Aza-c, CAS no 320-67-2), retinoic acid (RA, CAS no-302-79-4) and AM580 (CAS no- 102121-60-8) were purchased from Sigma Aldrich. The stock solution of all chemicals were prepared in high performance liquid chromatography (HPLC) grade dimethyl sulfoxide (DMSO) and stored at cool and dark place in 1ml amber glass vials with polytetrafluoroethylene-lined caps. The working solution were prepared in 1X embryo media (0.17 mM KCl, 0.33 mM CaCl_2_, 0.33 mM MgSO4 and 5 mM NaCl, pH 7). All chemicals and reagents used in the present study were of analytical grade.

### 2.3. Embryonic exposure

Viable embryos at 6-hour post fertilization (6hpf) or blastula stage were exposed under static condition in temperature-controlled incubator [VWR (Radnor, PA, USA), 28 ± 0.2 ^0^C] till 120 hpf. We first conducted initial range finding experiments with 0 - 200 μM IPP to identify mortality and morphological effects using Olympus SZ61 stereomicroscope equipped with Infinity 8 camera. 0.1% DMSO was used as negative control in the study. Embryos were exposed to 0 or 20 μM IPP for bioaccumulation and neurotransmitter studies but for LPR, exposure groups were nontoxic 0, 0.00002, 0.0002, 0.002, 0.02 and 0.2 μM IPP. For rest of the experiments, we choose two lower (0.02 and 0.2 μM) concentrations. For co-exposure studies, sublethal concentrations of 5-azacytidine (Aza-c, 7.81 μM), RA (1.9 pM) and AM580 (10 pM) were decided through range finding experiments and co-exposures were conducted with these concentrations in presence/absence of 0, 0.02 or 0.2 μM IPP.

### 2.4. Larval photomotor response

Experiments were set up in 96-well plates (Corning; one embryo and 100 μl test solution in a well), and treatment group containing 32 embryos (16 embryos/plate, total 2 plates) were exposed to different concentrations of IPP. The plates were covered with parafilm to avoid evaporation and wrapped with aluminum foil to prevent light. Larval photomotor response was measured using a Zebra box (Viewpoint, France) according to previous protocols (Dasgupta et al., 2023). Viewpoint analysis measures the photomotor response (every 6 sec) of 5-day old fish to alternate 3 min light and dark cycles, for 4 cycles each. Dead and deformed embryos were excluded and only 3^rd^ dark cycle was considered for analysis. Each exposure group contained 32 embryos.

### 2.5. RA signaling

To determine the effect of IPP on RA signaling, *in vitro* assay for 3 retinoic acid receptors (RARα, RARβ and RARγ) were conducted using Chinese hamster ovary (CHO) stably transfected with chimeric human RAR orthologues (Indigo Biosciences). The cells were exposed to 0.01 - 10 μM IPP and RA inhibitor (BMS195614 or CD2665). RA-induced luciferase activity was measured from mean luminescence values.

### 2.6. Whole mount Immunohistochemistry (IHC)

IHC was performed as described in Dasgupta et al. (2019) with slight modifications. After exposure, the embryos were fixed for 24 hrs in paraformaldehyde (4%) and kept in phosphate buffer saline (PBS, pH 7.4) at 4^0^C until they were processed further. Larvae were incubated in fresh bleaching solution (3% H_2_O_2_ and 0.5% KOH) for 30 min at room temperature and depigmentation was stopped by washing with 1X PBS. Larvae were then permeabilized in cold acetone by being incubated for 30 min at -20 ^0^C. Following this, embryos were kept in blocking buffer (1X PBST, 2% sheep serum and 2mg/ml bovine serum albumin) for 4 hrs and proceed with staining steps. For cytosine methylation, primary anti-5-methyl cytosine (1:200) and secondary IgG1λ (1:500) antibodies were used (Sigma Aldrich, St. Louis, MO). Ten embryos/treatment were imaged in ECHO Revolve R4 (San Diego, CA, USA) fluorescent microscope and Image J software was used to detect fluorescence intensity by measuring mean gray values.

### 2.7. Global DNA Methylation (whole embryo)

Genomic DNA (gDNA) was extracted from whole embryos (5 embryos/replicate, n = 10) using Zymo Research Quick DNA/RNA™ Mini preparation kit following manufacturer’s instructions. The embryos were homogenized in 1.5 ml RINO tubes (Next Advance) using DNA/RNA shield (1X, 600 μl) and 0.5 mm zirconium oxide beads in Bullet Blender Pro Series homogenizer (Next Advance, speed 8, time 5 min). The quantity and quality of gDNA was analyzed through Nanodrop Spectrophotometer (Thermo Scientific One). Global DNA methylation was determined calorimetric through MethylFlash Global DNA (5-mc) Elisa Easy kit (Epigentek) according to manufacturer’s instructions. The absorbance was measured at 450 nm using Synergy H1 hybrid plate reader (BioTek).

### 2.8. Global DNA Methylation (Eye and brain)

Eye and brain were harvested from whole embryo in Ringer solution (1.8 mM CaCl_2_, 2.9 mM KCl, 5 mM HEPES, 116 mM NaCl, pH 7.2) using Roboz surgical needles under Olympus SZ61 stereomicroscope as described by Zhang and Leung (2010). Ten eyes and five brains were used for each replicate (n = 4), washed with embryo media and homogenized in DNA/RNA lysis buffer for gDNA isolation. Level of DNA methylation was measured using same kit and protocol as for whole embryo global DNA methylation.

### 2.9. mRNA Sequencing

Embryos were exposed to solvent control (DMSO 0.02%) and IPP (0.2 μM) in 96 well plate and incubated at 28 ^0^C. Eyes and brains were dissected, collected in RINO tubes and homogenized in 300 μl RNAzol using Bullet Blender Pro Series homogenizer. Total RNA was extracted using Direct-Zol™ RNA miniprep kit according to the manufacturer’s instructions. RNA integrity was assessed with high sensitivity RNA screen tape in tape station (Agilent, 4200). Library preparation and sequencing was performed by Novogene (California, USA). The genes having adjusted p-values less than 0.05, were considered as differentially expressed. Gene Ontology (GO) enrichment analyses were performed on differentially expressed genes using SR plot, an online platform.

### 2.10. Statistical analysis

All statistical analyses were conducted, and graphs were generated using GraphPad Prism 9.0. For normalized data, one-way analysis of variance (ANOVA) followed by Dunnett’s posthoc test but for LPR assay Kruskal-Wallis followed by Dunn test were performed to find significance level at 0.01, 0.1 and 5 % (p < 0.001, p < 0.01 and p < 0.05, respectively). Paired-t test was used for neurotransmitter analysis.

## 3. Results

### 3.1. IPP exposures induce morphological abnormalities and hypoactive behavior in larvae

Embryonic exposure at 6hpf to higher concentrations of IPP (20 and 200 μM) showed phenotypic anomalies like pericardial edema, yolk sac edema and spinal curvature after 5dpf (**Fig. 1A**). The extent of pericardial edema increased significantly in exposed larvae compared to control (**Fig. 1B**). We then conducted larval photomotor response assay following IPP exposures at lower sublethal concentrations. Within this data, we found significant hypoactive locomotory behavior in exposed larvae at all tested concentrations. (**Fig. 1C**).

**Figure 1.**
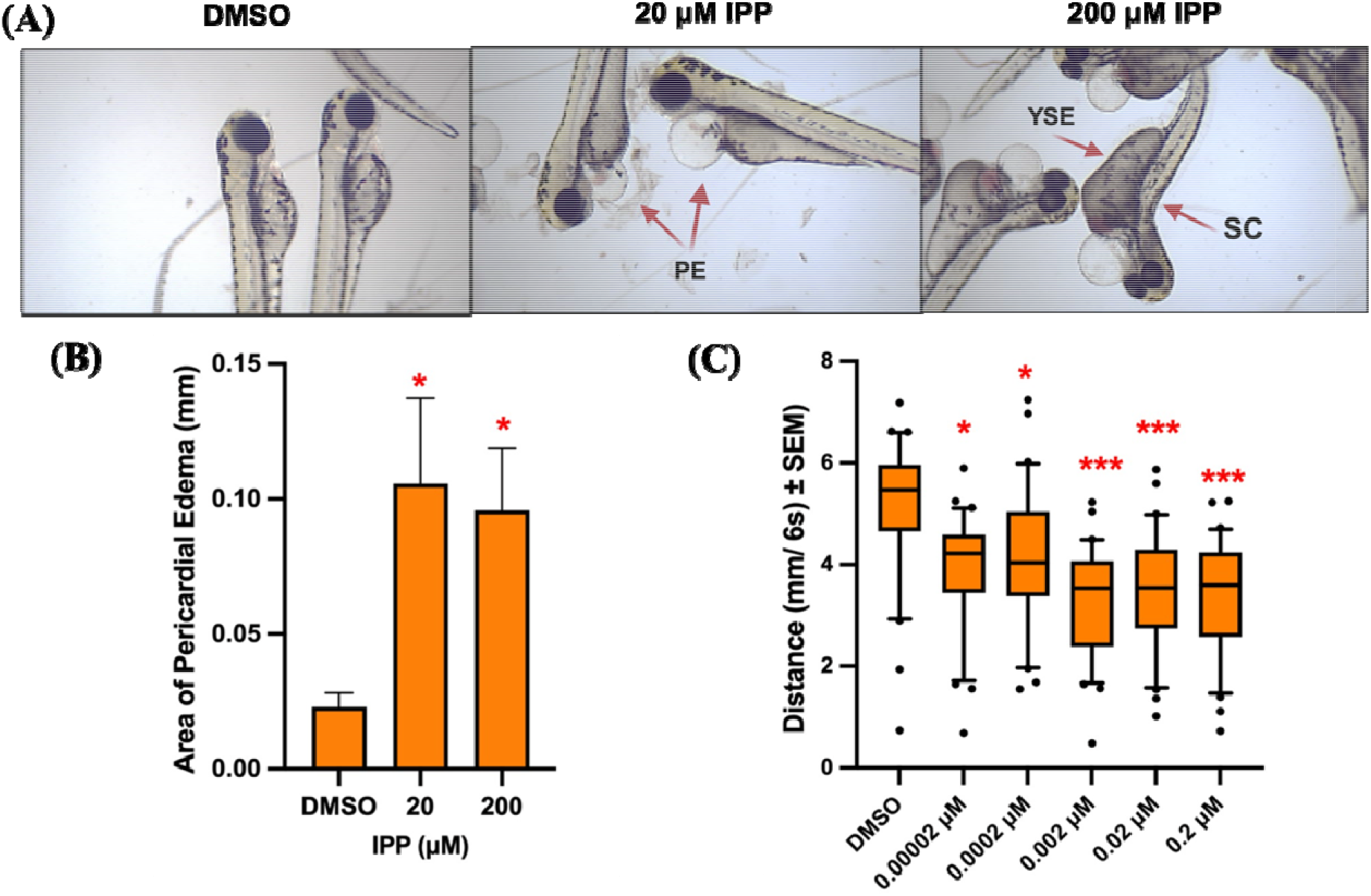
IPP induced phenotypic anomalies [pericardial edema (PE), yolk sac edema (YSE) and spinal curvature (SC)] (A), area of pericardial edema (B) and LPR assay (C) after 120 hpf For A and B, N=4, replicates of 10 embryos each in 24-well plates, **\***Statistically different from DMSO based on 1-way ANOVA followed by Dunnett’s test. For C, N=30-32 embryos/ exposure group, 96-well plates, **\***Statistically different from DMSO based Kruskal-Wallis followed by Dunn test

### 3.2. Reanalysis of prior mRNA sequencing data

Based on secondary analysis of our previous study (Dasgupta et al., 2021), we found that IPP exposure (20 μM) to zebrafish embryos (6hpf - 48hpf) significantly altered the expression DNA methyltransferase, and other genes related with neurotransmitter and retinoic acid signaling with in whole embryo (**Table 1**). IPP also affected various pathways related to eye development and visual perception (**Table 2**).

**Table 1.**
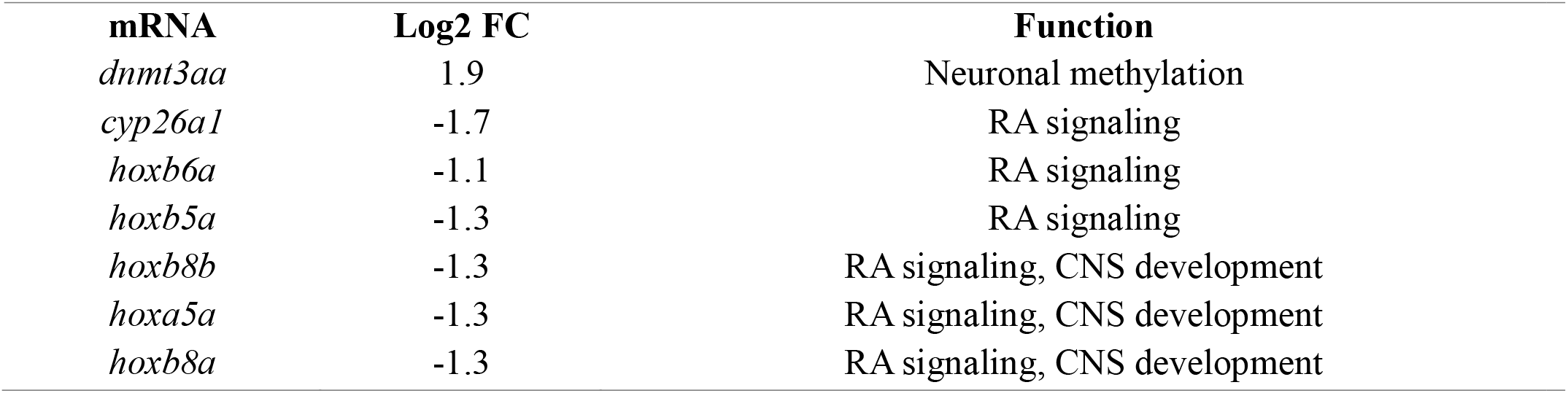
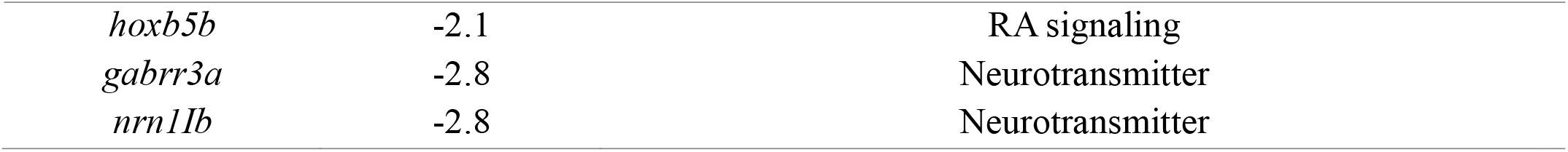
Differentially expressed genes and their functions when 6hpf embryos were exposed to 20 μM for 48 hpf.

**Table 2.**
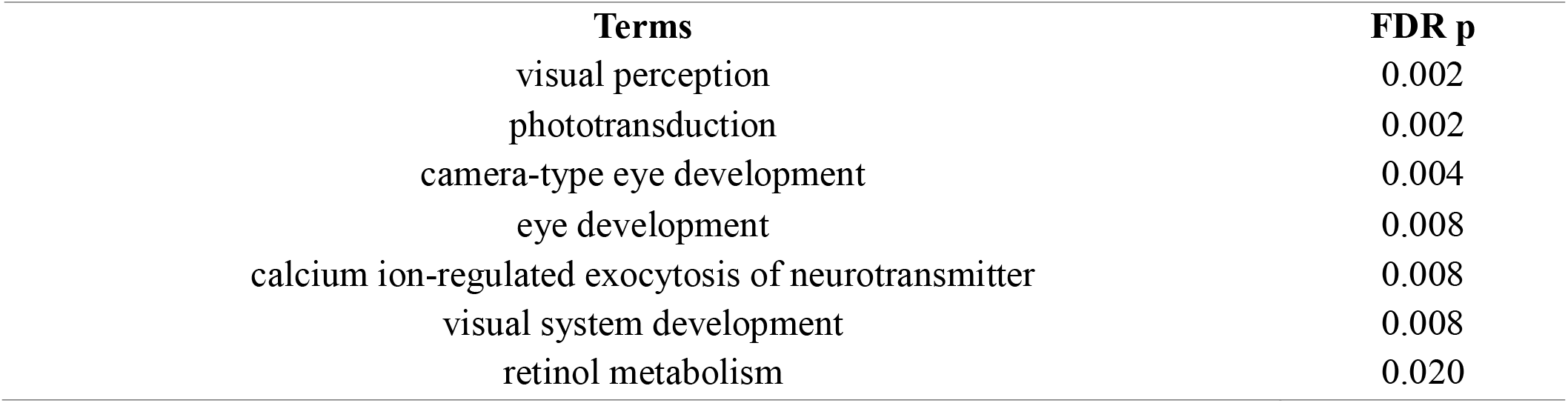
Gene ontology assessments of IPP induced mRNA disruptions.

### 3.3. IPP inhibits retinoic acid signaling

We conducted *in vitro* assays for retinoic acid signaling using CHO cells. In IPP-exposed cells, mean luminescence values declined significantly for retinoic acid receptor α (RARα) at 1 and 10 μM IPP **(Fig. 2A)**. We did not see any inhibitory effects on RARβ and RARγ in our study. Based on this, we hypothesized that IPP inhibits RA signaling and co-exposures with RA and AM580 (an RARα activator) will rescue the LPR phenotype. Contrary to our hypothesis we found that RA agonist were unable to rescue IPP induced hypoactivity **(Fig. 2B and 2C)**.

**Figure 2.**
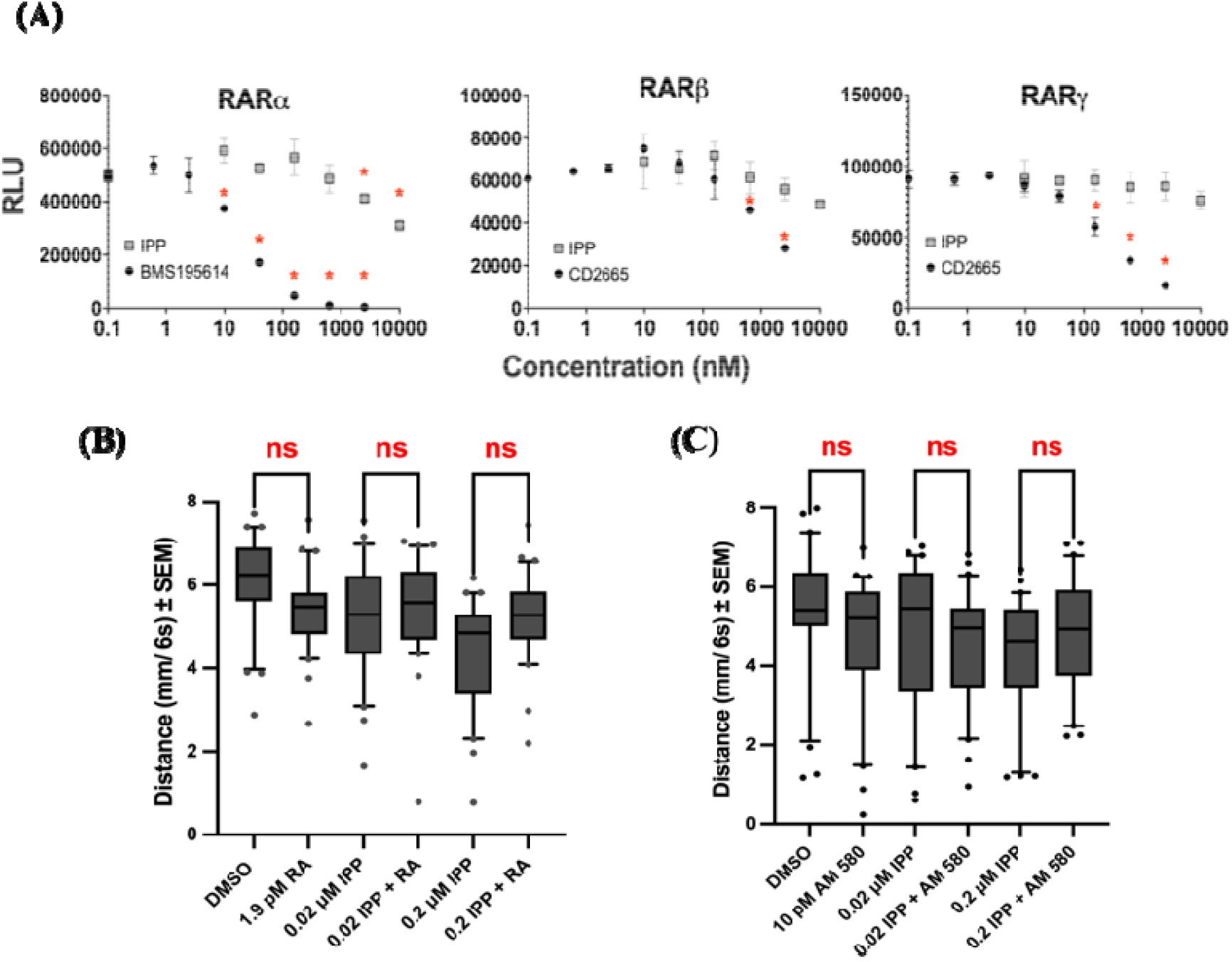
IPP inhibits RARα in Chinese hamster ovary (CHO) cells. Mean luminescence from RAR-expressing, RA activated CHO cells after exposure to various concentrations of IPP and retinoic acid inhibitor, BMS195614 or CD2665 (A), RLU= relative luminescence units. N=3. * p ≤ 0.05 with ANOVA, followed by Dunnett’s test. IPP-RA (B) and IPP-AM 580 (C) did not change IPP induced LPR responses N=30-32 embryos/ exposure group, 96-well plates. ns represents non-significant based Kruskal-Wallis followed by Dunn test

### 3.4. IPP induces DNA hypermethylation

To study DNA methylation, we first conducted IHC for 5-methylcytosine. Our data showed significant hypermethylation effects in head and whole body (**Fig. 3A, 3B and 3C**). We then co-exposed the embryos with Aza-c, a DNA methylation inhibitor (7.81 μM), to assess its mitigating role. We found that co-exposure significantly reversed IPP-induced hypermethylation in head and whole body (Fig. 3A and 3C). To complement this data, we also conducted an ELISA-based global DNA methylation assay; this also showed significant hypermethylation effects at both concentrations **(Fig. 3D)**. To validate whether DNA methylation is related with IPP induced hypoactivity, we conducted co-exposure LPR experiments with Aza-c and IPP. Our data shows that Aza-c showed significantly ameliorated IPP-induced effects, but also induced hyperactivity by itself. **(Fig. 3E)**.

**Figure 3.**
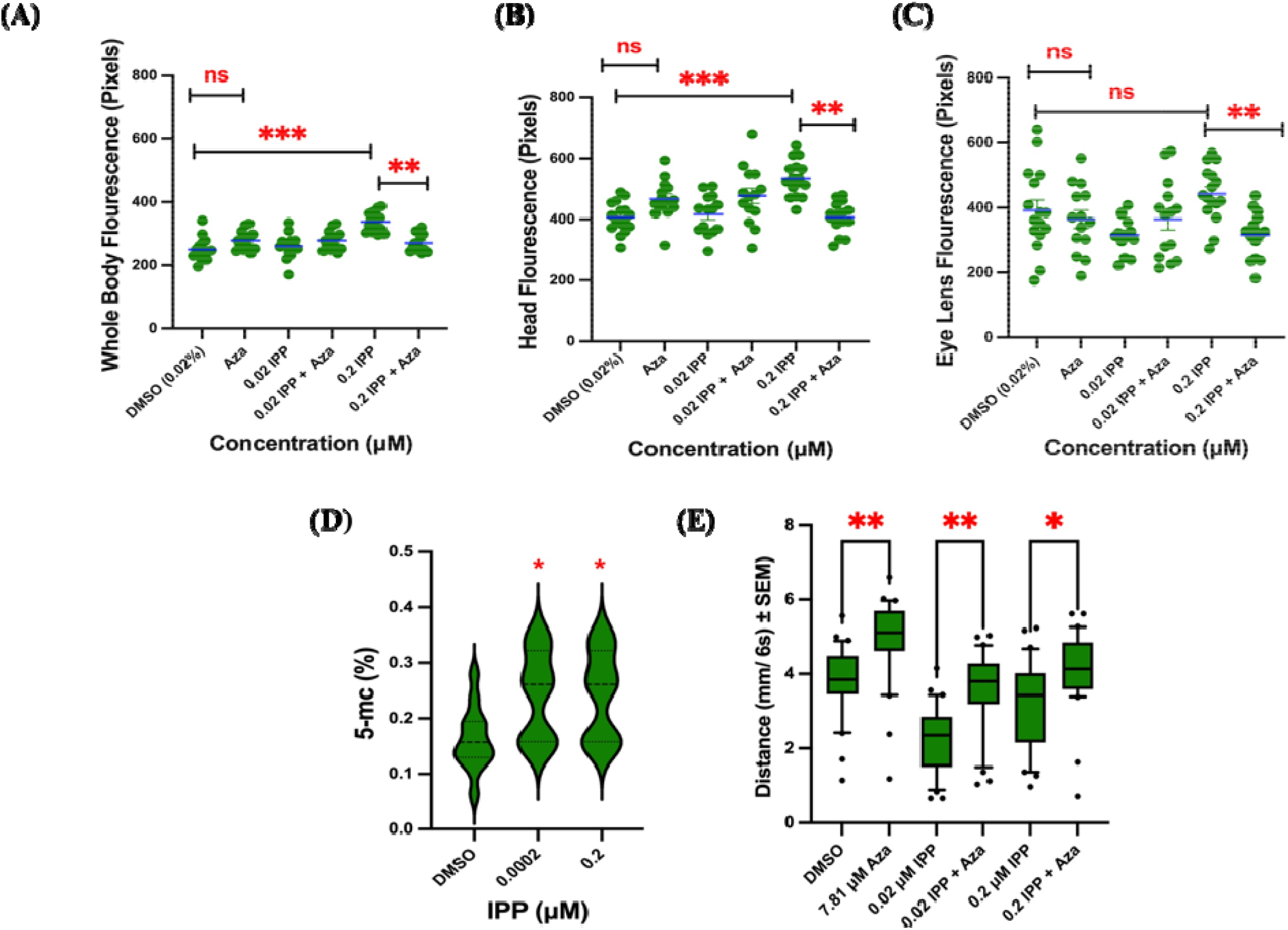
In situ 5-mc levels on co-exposure to IPP and Aza-c in whole body (A), head (B) and Eye lens (C), N=15 embryos/ replicate, 1-way ANOVA followed by Tukey’s test was used for statistics, ** (p ≤ 0.01) and *** (p ≤ 0.001) represent statistically different at 1% and 0.1 % significance level, respectively, ns represent no-significant difference. IPP induced global DNA hypermethylation (D), N=10, 5 embryos/ replicate, * represent statistically different from DMSO based on 1-way ANOVA followed by Dunnett’s test at p<0.05. IPP-Aza co-exposures differentially change LPR responses (E), N=30-32 embryos/ exposure group in 96-well plates. **\***Statistically different from DMSO based Kruskal-Wallis followed by Dunn test

### 3.5. IPP did not alter global DNA methylation in eyes and brain

Based on our prior whole embryonic sequencing data we then focused on impacts on eye and brain. We first conducted global DNA methylation studies; however, as seen in Fig 4, no significant differences were observed at both concentrations for these organs **(Fig. 4)**.

**Figure 4.**
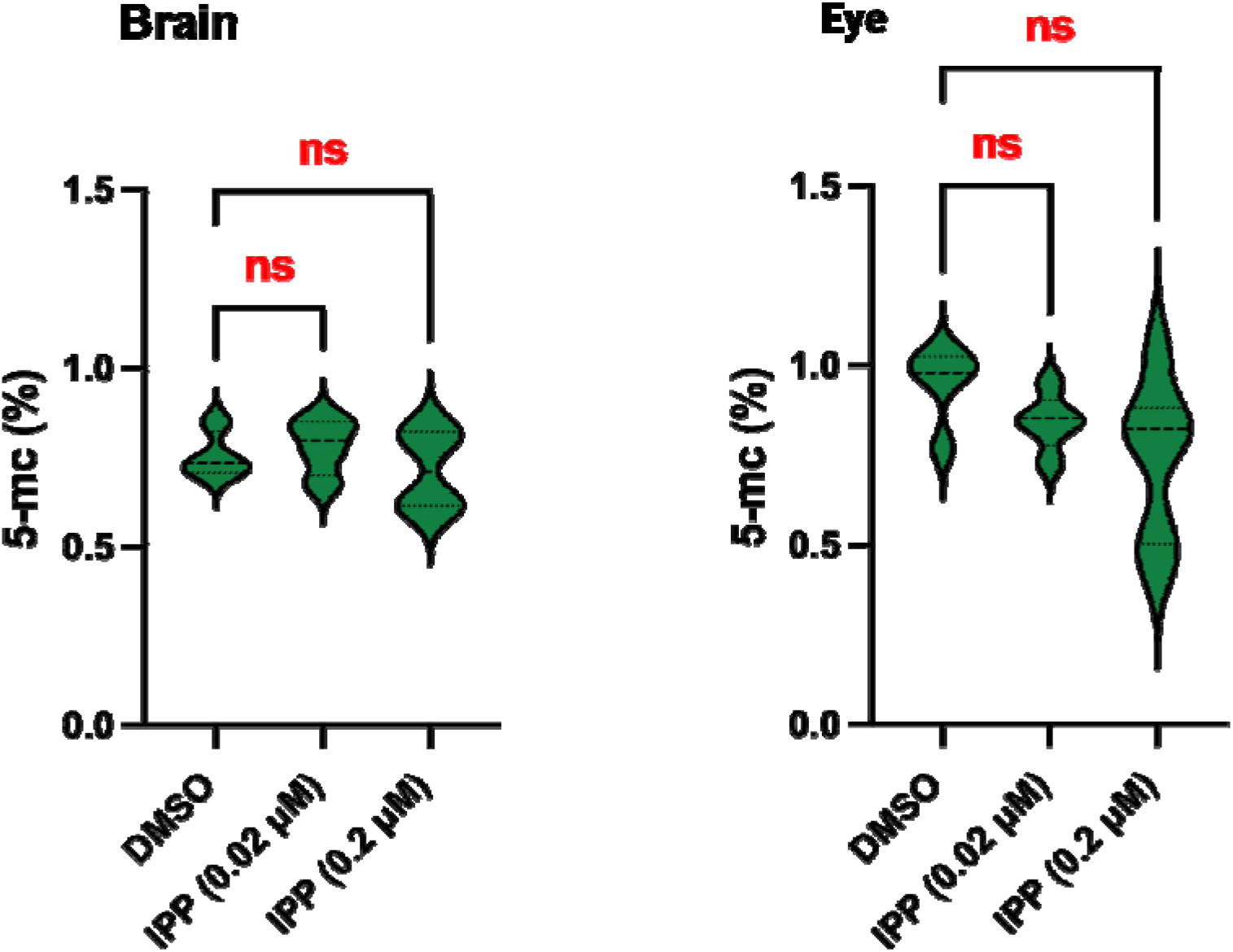
IPP did not show significant change in global DNA methylation from extracted brain and eye at 0.02 and 0.2 μM IPP. ns represents statistically non-significant based on 1-way ANOVA followed by Tukey’s test

### 3.6. mRNA sequencing shows IPP dysregulates genes related to eye and neurodevelopment

To examine genetic targets of IPP in brain and eye, we conducted mRNA-sequencing on extracted eyes and brain. Surprisingly, mRNA-seq data from brain at 0.2 μM IPP showed only 2 differentially expressed genes (DEGs); both were downregulated (data not shown). In contrast, our eye data showed 136 DEGs (41 upregulated and 94 downregulated) **(Fig. 5A and 5B)**. Notably, gene *batf2* (log_2_ FC = 7.30, p < 0.001) and *kcnip3b* (log_2_ FC = -1.44, p < 0.001) showed the highest level of differential expression. We also observed decreased mRNA levels of DNA (cytosine-5-)-methyltransferase 3 beta, duplicate a (*dnmt3ba*) in our study. Gene ontology revealed that these DEGs were associated with various biological and molecular pathways, but the top impacted pathways were, activity of voltage gated ion channels, synaptic transmission and neurotransmitter signaling **(5C and 5D)**.

**Figure 5.**
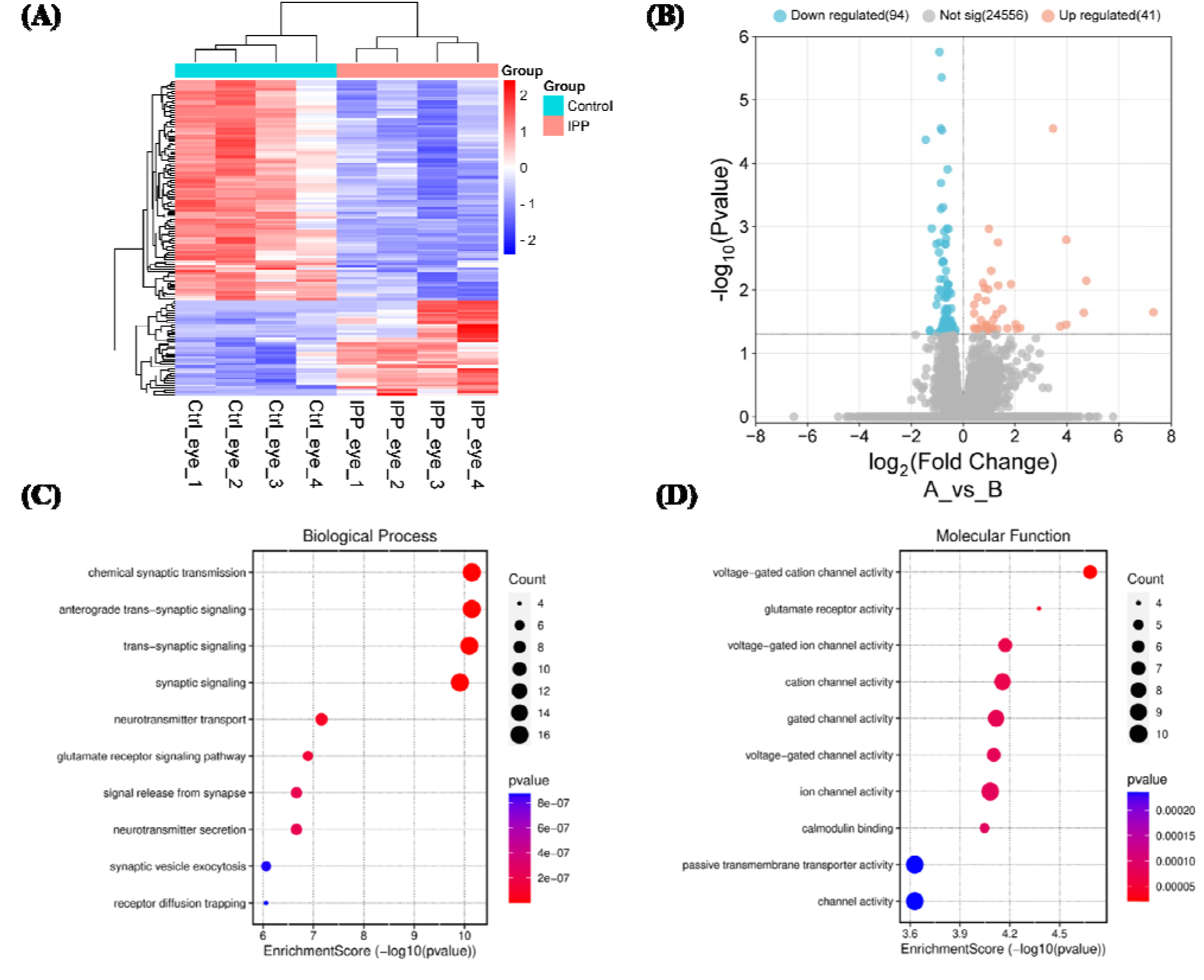
Heat map (A) and volcano plot (B) of DEG in eyes, when embryos were exposed to 0.2 μM IPP. Gene ontology reveals that IPP affects various biological (C) and molecular (D) pathways in developing eyes.

## 4. Discussion

Extensive work has been done on toxicity of different FRs, but very limited information is available on IPP as an isomeric mixture. In this study, we used zebrafish embryos to evaluate the developmental effects of IPP. We specifically focused on larval photomotor response and leveraged this assay as a readout for potential neurotoxic effects. Zebrafish exhibit sensitive nervous system and range of behaviors (learning, memory and locomotion) which are essential to assess the impact of toxicants on development. High throughput behavioral assays with real time tracking using larval zebrafish have become a standard technique nowadays in toxicological research (Bailey et a., 2013; Basnet et al., 2019). Using this method provide valuable insights into the functioning dynamics of the stress circuitry, integrity of nervous system and possible risk for disorders (McAtee and Abdelmoneim, 2024). Prior studies have shown that this assay can be driven by combined output of multisensory input and the coordination of skeletal and neuromuscular signals (Huang et al., 2022; Wang et al., 2022). Significant hypoactive behavior that we observed at nontoxic IPP concentrations can potentially be an indicator of neurodevelopmental toxicity. Indeed, components of IPP have been shown to elicit differential behavioral responses; for example, Zhang et al. (2023) observed decreased swimming speed in zebrafish embryos exposed to TPP for 120hpf.

We also interrogated the role of RA signaling, since our *in vitro* assessments showed RAR inhibition α. RA signaling has important roles in neurodevelopmental processes by regulating gene expression that can be impaired by toxicants (Chao et al., 2021; Menegola et al., 2021). RARα inhibition and non-mitigating role of RA agonists in our study suggest that neurobehavioral outcomes may involve complex mechanism beyond simple receptor activation. Xu et al. (2013) reported that acute exposure to PBDEs declined retinoid profile and downregulated expression of retinol dehydrogenase (*raldh2*), RA binding proteins (*crabp1a* and *crabp2a*) and RAR subunit (*raraa*) in zebrafish. Consistent with our results, Hawkey et al. (2024) observed that developmental exposure to environmental toxicants interfere with RA synthesis and catabolic genes which contribute for behavioral disruptions in zebrafish larvae. Inhibitory effects of IPP on RA signaling might impact the development of retinal photoreceptor and eye morphogenesis resulted in visual impairment (Xu et al., 2013). These findings highlight the potential role of RAR α in mediating the neurotoxic effects of IPP and reflect that other factors or pathways may be involved in IPP induced hypoactivity.

DNA methylation is a common epigenetic modification that affect gene activity and cellular function during development. In general, CpG site modification have a significant impact on binding of transcription factors and gene expression (Jones, 2012). Based on prior whole embryo RNA seq results (Dasgupta et al., 2021), we focused on DNA methylation as a potential IPP target and conducted 5-mc based assays. The observed hypermethylation effects in response to IPP exposure can be related with upregulation of *dnmt3ba* in our study. According to Moore et al. (2013), hypomethylation promotes gene transcription while hypermethylation results in gene silencing, this can be linked with downregulation of neurodevelopmental genes in our previous study (Dasgupta et al., 2021). Tris (1, 3-dichloro-2 propyl) phosphate (TDCIPP), another OPFR have also been observed to increase cytosine methylation in yolk sac and cell mass region of zebrafish embryos during blastula and gastrula stage, respectively (Avila-Barnard et al., 2023). Olsvik et al. (2019) observed that bisphenol A exposure to zebrafish embryos altered the methylation patterns of genes associated with neurodevelopment resulted in abnormal swimming. Further studies, including targeted analyses of gene expression and functional assays, are needed to fully elucidate the consequences of DNA hypermethylation on neurodevelopment.

For organ specific toxicity, we performed global DNA methylation and mRNA seq on dissected brain and eye. Non-significant changes in global DNA methylation of brains and eyes suggest that the tissues may have distinct epigenetic responses, or it is worth considering that global DNA methylation assay may not be sensitive enough to detect subtle alterations in these organs. In brain, we noticed downregulation of somatolactin beta (Slβ, log2 FC -2.77)), a multifunctional gene having role in pigmentation, energy balance, stress response, growth and development. Liu et al. (2018) related the alteration in transcript levels of Slα and Slβ with endocrine disruption in rare minnow on exposure to cadmium. Gene ontology analyses revealed that IPP impacts eye development by affecting phototransduction, voltage gated channels, synaptic transmission and neurotransmitter signaling. These alterations can impact the ability of the eye to detect and respond to light stimuli, functioning of neurons in retina for the generation and propagation of electric signals and proper neuronal communication. Disruption of these pathways may lead to aberrant neuronal firing patterns and impaired muscle function, ultimately affecting swimming behavior. It is assumed that these alterations may disrupt the finely tuned balance of neurotransmitters and calcium ions, leading to impaired muscle contraction and coordination. Shi et al. (2018, 2019) reported that TPHP exposure altered the levels of visual and muscular proteins that may be responsible for lethargic swimming in zebrafish larvae. Through behavioral assay, RA signaling and transcriptome analysis we demonstrated that IPP at sublethal concentrations caused developmental neurotoxicity.

## 5. Conclusions

Our study indicates that IPP induced developmental toxicity, within zebrafish embryos through morphological abnormalities and disrupting larval photo motor response. We conclude that IPP potentially affect DNA methylation, neurotransmitter levels, synaptic signaling, muscle contraction and photoreceptor system which impaired neuronal development, body coordination and vision that would be the probable reason for hypoactive swimming behavior. These studies are important to evaluate the toxic potential of environmental chemicals and determination of toxicological endpoints for regulatory purposes.

## 6. Acknowledgements

This research was made possible with startup funds from Clemson University, College of Science and Department of Biological Sciences. We acknowledge Mr. John Smink, Facility Manager, Aquatic Animal Research Laboratory for husbandry of zebrafish and Dr. Robyn Tanguay (Oregon State University) for generously providing us with 5D founder fish. We also acknowledge Clemson University Genomics and Bioinformatics Facility for use of Tape station and Nanodrop.

## Notes

### Competing Interest Statement

The authors have declared no competing interest.

## References

Araki, A., Bastiaensen, M., Bamai, Y.A., Van den Eede, N., Kawai, T., Tsuboi, T., Ketema, R.M., Covaci, A. and Kishi, R., 2018. Associations between allergic symptoms and phosphate flame retardants in dust and their urinary metabolites among school children. Environment international, 119, pp.438–446.

Australian Government (Department of Health and Aged Care), Evaluation Statement., 2023. Triphenyl phosphate and diphenyl phosphate.

Avila-Barnard, S., Dasgupta, S., Cheng, V., Reddam, A., Wiegand, J.L. and Volz, D.C., 2022. Tris (1, 3-dichloro-2-propyl) phosphate disrupts the trajectory of cytosine methylation within developing zebrafish embryos. Environmental research, 211, p.113078.

Bailey, J., Oliveri, A. and Levin, E.D., 2013. Zebrafish model systems for developmental neurobehavioral toxicology. Birth Defects Research Part C: Embryo Today: Reviews, 99(1), pp.14–23.

Banks, A.P.W., Engelsman, M., He, C., Wang, X. and Mueller, J.F., 2020. The occurrence of PAHs and flame-retardants in air and dust from Australian fire stations. Journal of Occupational and Environmental Hygiene, 17(2-3), pp.73–84.

Basnet, R.M., Zizioli, D., Taweedet, S., Finazzi, D. and Memo, M., 2019. Zebrafish larvae as a behavioral model in neuropharmacology. Biomedicines, 7(1), p.23.

Butt, C.M., Congleton, J., Hoffman, K., Fang, M. and Stapleton, H.M., 2014. Metabolites of organophosphate flame retardants and 2-ethylhexyl tetrabromobenzoate in urine from paired mothers and toddlers. Environmental science & technology, 48(17), pp.10432–10438.

Butt, C.M., Hoffman, K., Chen, A., Lorenzo, A., Congleton, J. and Stapleton, H.M., 2016. Regional comparison of organophosphate flame retardant (PFR) urinary metabolites and tetrabromobenzoic acid (TBBA) in mother-toddler pairs from California and New Jersey. Environment international, 94, pp.627–634.

Castorina, R., Bradman, A., Stapleton, H.M., Butt, C., Avery, D., Harley, K.G., Gunier, R.B., Holland, N. and Eskenazi, B., 2017. Current-use flame retardants: Maternal exposure and neurodevelopment in children of the CHAMACOS cohort. Chemosphere, 189, pp.574–580.

Chen, G., Jin, Y., Wu, Y., Liu, L. and Fu, Z., 2015. Exposure of male mice to two kinds of organophosphate flame retardants (OPFRs) induced oxidative stress and endocrine disruption. Environmental toxicology and pharmacology, 40(1), pp.310–318.

Cho, K., Lee, S.M., Heo, J., Kwon, Y.M., Chung, D., Yu, W.J., Bae, S.S., Choi, G., Lee, D.S. and Kim, Y., 2021. Retinaldehyde dehydrogenase inhibition-related adverse outcome pathway: Potential risk of retinoic acid synthesis inhibition during embryogenesis. Toxins, 13(11), p.739.

Dasgupta, S., Cheng, V., Vliet, S.M., Mitchell, C.A. and Volz, D.C., 2018. Tris (1, 3-dichloro-2-propyl) phosphate exposure during the early-blastula stage alters the normal trajectory of zebrafish embryogenesis. Environmental science & technology, 52(18), pp.10820–10828.

Dasgupta, S., Dunham, C.L., Truong, L., Simonich, M.T., Sullivan, C.M. and Tanguay, R.L., 2021. Phenotypically anchored mRNA and miRNA expression profiling in zebrafish reveals flame retardant chemical toxicity networks. Frontiers in Cell and Developmental Biology, 9, p.663032.

Dasgupta, S., LaDu, J.K., Garcia, G.R., Li, S., Tomono-Duval, K., Rericha, Y., Huang, L. and Tanguay, R.L., 2023. A CRISPR-Cas9 mutation in sox9b long intergenic noncoding RNA (slincR) affects zebrafish development, behavior, and regeneration. Toxicological Sciences, 194(2), pp.153–166.

Dasgupta, S., Vliet, S.M., Cheng, V., Mitchell, C.A., Kirkwood, J., Vollaro, A., Hur, M., Mehdizadeh, C. and Volz, D.C., 2019. Complex interplay among nuclear receptor ligands, cytosine methylation, and the metabolome in driving tris (1, 3-dichloro-2-propyl) phosphate-induced epiboly defects in zebrafish. Environmental science & technology, 53(17), pp.10497–10505.

Daston, G., Faustman, E., Ginsberg, G., Fenner-Crisp, P., Olin, S., Sonawane, B., Bruckner, J., Breslin, W. and McLaughlin, T.J., 2004. A framework for assessing risks to children from exposure to environmental agents. Environmental health perspectives, 112(2), pp.238–256.

Ekpe, O.D., Choo, G., Barceló, D. and Oh, J.E., 2020. Introduction of emerging halogenated flame retardants in the environment. In Comprehensive analytical chemistry (Vol. 88, pp. 1–39). Elsevier.

Fu, Z., Xie, H.B., Elm, J., Liu, Y., Fu, Z. and Chen, J., 2021. Atmospheric autoxidation of organophosphate esters. Environmental Science & Technology, 56(11), pp.6944–6955.

Guo, X., Mu, T., Xian, Y., Luo, D. and Wang, C., 2016. Ultra-performance liquid chromatography tandem mass spectrometry for the rapid simultaneous analysis of nine organophosphate esters in milk powder. Food Chemistry, 196, pp.673–681.

Han, X., Hao, Y., Li, Y., Yang, R., Wang, P., Zhang, G., Zhang, Q. and Jiang, G., 2020. Occurrence and distribution of organophosphate esters in the air and soils of Ny-Ålesund and London Island, Svalbard, Arctic. Environmental pollution, 263, p.114495.

Harrad, S., Brommer, S. and Mueller, J.F., 2016. Concentrations of organophosphate flame retardants in dust from cars, homes, and offices: An international comparison. Emerging Contaminants, 2(2), pp.66–72.

Hawkey, A.B., Shekey, N., Dean, C., Asrat, H., Koburov, R., Holloway, Z.R., Kullman, S.W. and Levin, E.D., 2024. Developmental exposure to pesticides that disrupt retinoic acid signaling causes persistent retinoid and behavioral dysfunction in zebrafish. Toxicological Sciences, p.kfae001.

He, H., Gao, Z., Zhu, D., Guo, J., Yang, S., Li, S., Zhang, L. and Sun, C., 2017. Assessing bioaccessibility and bioavailability of chlorinated organophosphorus flame retardants in sediments. Chemosphere, 189, pp.239–246.

Hoffman, K., Butt, C.M., Chen, A., Limkakeng Jr, A.T. and Stapleton, H.M., 2015. High exposure to organophosphate flame retardants in infants: associations with baby products. Environmental science & technology, 49(24), pp.14554–14559.

Hoffman, K., Lorenzo, A., Butt, C.M., Adair, L., Herring, A.H., Stapleton, H.M. and Daniels, J.L., 2017. Predictors of urinary flame-retardant concentration among pregnant women. Environment international, 98, pp.96–101.

Huang, W., Xiao, J., Shi, X., Zheng, S., Li, H., Liu, C. and Wu, K., 2022. Effects of di-(2-ethylhexyl) phthalate (DEHP) on behavior and dopamine signaling in zebrafish (Danio rerio). Environmental Toxicology and Pharmacology, 93, p.103885.

Huang, Y., Tan, H., Li, L., Yang, L., Sun, F., Li, J., Gong, X. and Chen, D., 2020. A broad range of organophosphate tri-and di-esters in house dust from Adelaide, South Australia: Concentrations, compositions, and human exposure risks. Environment International, 142, p.105872.

Jones, P.A., 2012. Functions of DNA methylation: islands, start sites, gene bodies and beyond. Nature reviews genetics, 13(7), pp.484–492.

Liu, X.H., Xie, B.W., Wang, Z.J. and Zhang, Y.G., 2018. Characterization and expression analyses of somatolactin-α and-β genes in rare minnows (Gobiocypris rarus) following waterborne cadmium exposure. Fish physiology and biochemistry, 44, pp.983–995.

Mcatee, D. and Abdelmoneim, A., 2024. Effects of developmental exposure to arsenic species on behavioral stress responses in larval zebrafish and implications for stress-related disorders. Toxicological Sciences, p.kfae074.

Menegola, E., Veltman, C.H., Battistoni, M., Di Renzo, F., Moretto, A., Metruccio, F., Beronius, A., Zilliacus, J., Kyriakopoulou, K., Spyropoulou, A. and Machera, K., 2021. An adverse outcome pathway on the disruption of retinoic acid metabolism leading to developmental craniofacial defects. Toxicology, 458, p.152843.

Mihajlović, I., Miloradov, M.V. and Fries, E., 2011. Application of Twisselmann extraction, SPME, and GC-MS to assess input sources for organophosphate esters into soil. Environmental science & technology, 45(6), pp.2264–2269.

Moore, L.D., Le, T. and Fan, G., 2013. DNA methylation and its basic function. Neuropsychopharmacology, 38(1), pp.23–38.

Olsvik, P.A., Whatmore, P., Penglase, S.J., Skjærven, K.H., Anglès d’Auriac, M. and Ellingsen, S., 2019. Associations between behavioral effects of bisphenol A and DNA methylation in zebrafish embryos. Frontiers in Genetics, 10, p.184.

Ospina, M., Jayatilaka, N.K., Wong, L.Y., Restrepo, P. and Calafat, A.M., 2018. Exposure to organophosphate flame retardant chemicals in the US general population: Data from the 2013–2014 National Health and Nutrition Examination Survey. Environment international, 110, pp.32–41.

Rice, D. and Barone Jr, S., 2000. Critical periods of vulnerability for the developing nervous system: evidence from humans and animal models. Environmental health perspectives, 108(suppl 3), pp.511–533.

Shi, Q., Tsui, M.M., Hu, C., Lam, J.C., Zhou, B. and Chen, L., 2019. Acute exposure to triphenyl phosphate (TPhP) disturbs ocular development and muscular organization in zebrafish larvae. Ecotoxicology and Environmental Safety, 179, pp.119–126.

Shi, Q., Wang, M., Shi, F., Yang, L., Guo, Y., Feng, C., Liu, J. and Zhou, B., 2018. Developmental neurotoxicity of triphenyl phosphate in zebrafish larvae. Aquatic toxicology, 203, pp.80–87.

Tachachartvanich, P., Rusit, X., Tong, J., Mann, C. and La Merrill, M.A., 2024. Perinatal triphenyl phosphate exposure induces metabolic dysfunctions through the EGFR/ERK/AKT signaling pathway: Mechanistic in vitro and in vivo studies. Ecotoxicology and Environmental Safety, 269, p.115756.

Volz, D.C., Leet, J.K., Chen, A., Stapleton, H.M., Katiyar, N., Kaundal, R., Yu, Y. and Wang, Y., 2016. Tris (1, 3-dichloro-2-propyl) phosphate induces genome-wide hypomethylation within early zebrafish embryos. Environmental science & technology, 50(18), pp.10255–10263.

Wade, M.G., Kawata, A., Rigden, M., Caldwell, D. and Holloway, A.C., 2019. Toxicity of flame retardant isopropylated triphenyl phosphate: liver, adrenal, and metabolic effects. International journal of toxicology, 38(4), pp.279–290.

Wang, X., Shi, X., Zheng, S., Zhang, Q., Peng, J., Tan, W. and Wu, K., 2022. Perfluorooctane sulfonic acid (PFOS) exposures interfere with behaviors and transcription of genes on nervous and muscle system in zebrafish embryos. Science of The Total Environment, 848, p.157816.

Wei, G.L., Li, D.Q., Zhuo, M.N., Liao, Y.S., Xie, Z.Y., Guo, T.L., Li, J.J., Zhang, S.Y. and Liang, Z.Q., 2015. Organophosphorus flame retardants and plasticizers: sources, occurrence, toxicity and human exposure. Environmental pollution, 196, pp.29–46.

Witchey, S.K., Sutherland, V., Collins, B., Roberts, G., Shockley, K.R., Vallant, M., Krause, J., Cunny, H., Waidyanatha, S., Mylchreest, E. and Sparrow, B., 2023. Reproductive and developmental toxicity following exposure to organophosphate ester flame retardants and plasticizers, triphenyl phosphate and isopropylated phenyl phosphate, in Sprague Dawley rats. Toxicological Sciences, 191(2), pp.374–386.

Xie, J., Tu, H., Chen, Y., Chen, Z., Yang, Z. and Liu, Y., 2023. Triphenyl phosphate induces clastogenic effects potently in mammalian cells, human CYP1A2 and 2E1 being major activating enzymes. Chemico-Biological Interactions, 369, p.110259.

Xing, L., Tao, M., Zhang, Q., Kong, M., Sun, J., Jia, S. and Liu, C.H., 2020. Occurrence, spatial distribution and risk assessment of organophosphate esters in surface water from the lower Yangtze River Basin. Science of the Total Environment, 734, p.139380.

Xu, T., Chen, L., Hu, C. and Zhou, B., 2013. Effects of acute exposure to polybrominated diphenyl ethers on retinoid signaling in zebrafish larvae. Environmental Toxicology and Pharmacology, 35(1), pp.13–20.

Yao, C., Yang, H. and Li, Y., 2021. A review on organophosphate flame retardants in the environment: Occurrence, accumulation, metabolism and toxicity. Science of the Total Environment, 795, p.148837.

Zhang, L. and Leung, Y.F., 2010. Microdissection of zebrafish embryonic eye tissues. JoVE (Journal of Visualized Experiments), (40), p.e2028.

Zhang, Q., Zheng, S., Shi, X., Luo, C., Huang, W., Lin, H., Peng, J., Tan, W. and Wu, K., 2023. Neurodevelopmental toxicity of organophosphate flame retardant triphenyl phosphate (TPhP) on zebrafish (Danio rerio) at different life stages. Environment International, 172, p.107745.

